# Unbiased clade age estimation using a Bayesian Brownian Bridge

**DOI:** 10.1101/2021.04.03.438104

**Authors:** Daniele Silvestro, Christine D. Bacon, Wenna Ding, Qiuyue Zhang, Philip C. J. Donoghue, Alexandre Antonelli, Yaowu Xing

## Abstract

In a recent paper^1^ we presented a new model, the Bayesian Brownian Bridge (BBB), to infer clade age based on fossil evidence and modern diversity. We benchmarked the method with extensive simulations, including a wide range of diversification histories and sampling heterogeneities that go well beyond the necessarily simplistic model assumptions. Applying BBB to 198 angiosperm families, we found that their fossil record is compatible with clade origins earlier than most contemporary palaeobotanical interpretations. In particular, we estimated with high probability that crown-angiosperms originated before the Cretaceous (> 145 Ma). Budd and colleagues^2^ critique our study, arguing that the BBB model is biased towards older estimates when fossil data are scarce or absent, that our underlying fossil dataset is unsound, that our clade age estimates are therefore biased by early diverging lineages that are underrepresented in the fossil record, and that pooling of fossil data for analysis at higher taxonomic ranks overcomes these biases. Here, we explore their points and perform new simulations to show that their critique has no merit.

---

Budd and colleagues speculate that, if there were no fossils at all, our model would be guaranteed (P = 1 – 0.483^198^) to find support for a pre-Cretaceous origin of any set of 198 clades, because it would sample root ages from the prior (here set to a uniform distribution *U*[0, 300]). This is false. To demonstrate this, we simulated 200 datasets with root ages randomly drawn from a uniform distribution *U*[180, 10] (i.e. the settings used in our published simulations) and without any fossils. BBB analysis of these extant-only datasets yielded 200 clade estimates (posterior mean) none of which were older than 66 Ma and only in seven datasets did the 95% CI reach into the Jurassic (> 145 Ma) – despite simulations including 47 clades of Jurassic origin (Extended Data Table 1). The combined probability that all 200 clades originated in the Cretaceous or later was 0.84 (i.e., 63 orders of magnitude higher than the probability predicted by Budd and colleagues). This is comparable to (but slightly more conservative than) the prior probability (P = 0.483) they indicate as appropriate. Thus, in the absence of fossil data, the BBB model might be biased but towards younger ages – not older, which undermines Budd and colleagues’ central claim.

Budd and colleagues simulate an entirely unrealistic scenario in which 198 clades all originate at the same time (120 Ma) and then diversify under speciation and extinction rates that are constant through time and identical across all clades. In contrast, overwhelming evidence from palaeontological and phylogenetic research demonstrate widespread and substantial heterogeneity in speciation and extinction rates across the tree of life in general, and specifically in the evolution of angiosperm clades^3-6^. Hence, their simulations reflect an aberrant case, in which the BBB model does not provide the correct answer. To demonstrate this, we simulated 198 clades rooted at 120 Ma under a more realistic Brownian bridge, which, unlike constant-rate birth-death models, allows for heterogeneities in the diversification process. 182 clades (92%) were correctly inferred to originate in the Cretaceous (16 older, two younger; Fig. 1a; Extended Data Table 2). In 160 clades (81%) the estimated ages were correctly assigned to the early Cretaceous with 95.5% coverage (fraction of estimates with 95% CI overlapping the true root age). These results differ markedly from those of Budd and colleagues^2^ (their Fig. 1C), highlighting the robustness of our model and demonstrating that their claims of biases in our model are artefacts of their own flawed simulation strategy.

**Figure 1.**
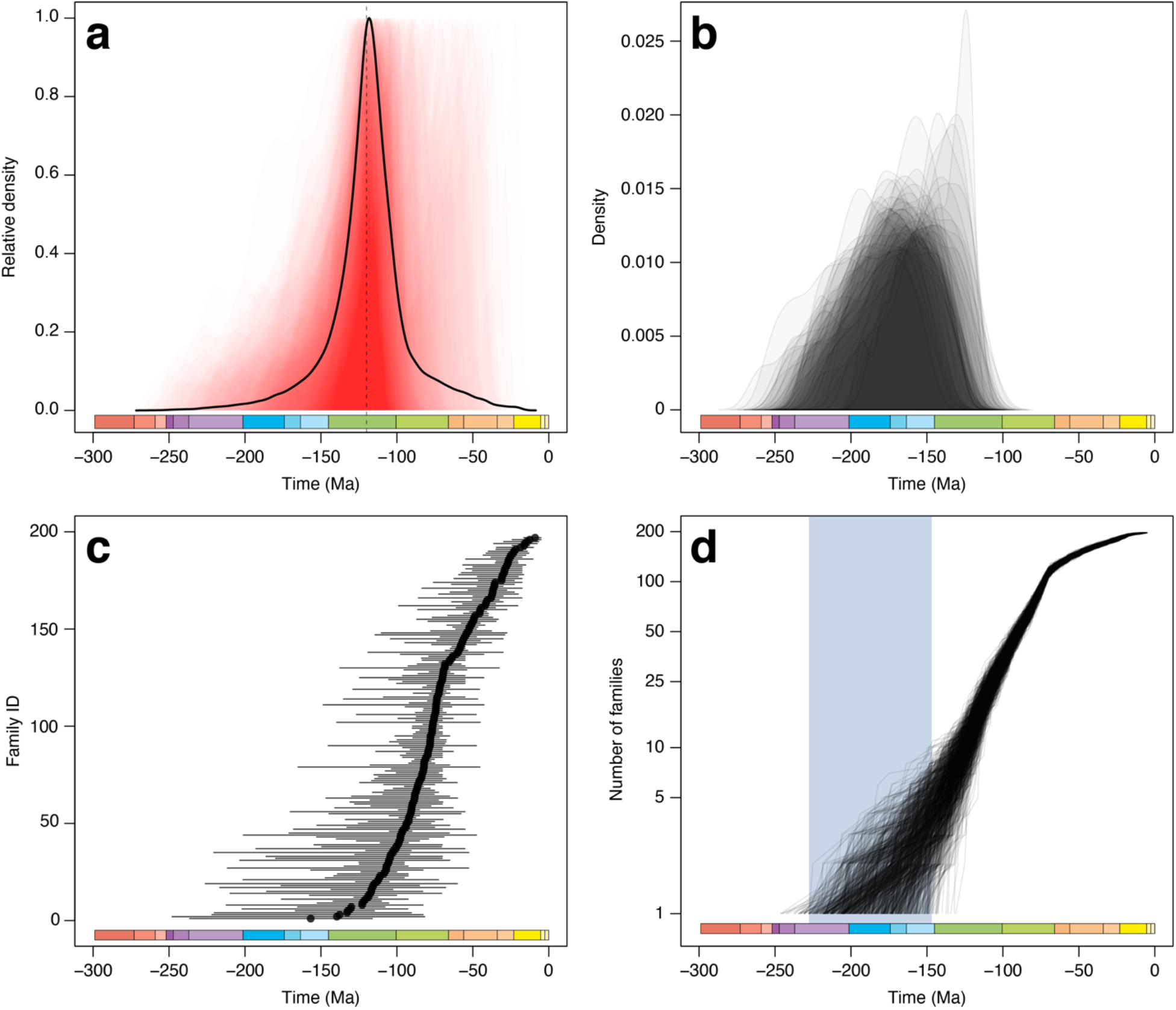
**a**) Estimated ages from 198 simulated clades with true time of origin set to 120 Ma (reflecting the simulations suggested by Budd et al.^2^). Red shaded areas show the individual posterior distributions, while the black line shows the density of the combined clade ages across all simulations. These results show that the BBB model returns unbiased results as shown in our previous simulations^1^ and that the coverage of our estimates is high (95.5% in this example; full results are available in Extended Data Table 2). b) Estimated crown ages from 50 simulations each of which consisted of 198 clades. In each simulation the ages of the simulated 198 subclades were distributed according to a birth-death process with time of origin set to 120 Ma. In 94% of the simulations a Cretaceous crown age was (correctly) included in the estimated 95% credible interval (full results are available in Extended Data Table 3). **c)** Analysis of the angiosperm fossil record at the family level after removing the records considered problematic by Budd et al.^2^ (detailed results are available in Extended Data Table 5). Circles represent posterior mean estimates of the clade ages while the horizontal bars show the 95% credible intervals. These results are highly consistent with our previous estimates^1^. **d)** lineage accumulation through time based on the estimated ages of the families shown in c). The shaded blue area shows the estimated 95% interval for the age of crown-angiosperms based on the ages of the sampled families. The interval spans from the Triassic to the late Jurassic, consistently with our previous estimates, with an estimated probability of a pre-Cretaceous origin of angiosperms equal to P = 0.986.

Paradoxically, Budd and colleagues’ simulation assumption, that 198 clades all originated at the same time, is incompatible with the very birth-death process they then use to simulate the diversity trajectory within each clade (where, by construction, the probability of more than 1 branching event at a given time is 0). It is illogical to expect the birth-death process to apply to diversification within clades but not to the origination of the clades themselves. To address this, we simulated 50 x 198 clade datasets in which the clade ages are distributed according to a birth-death process starting at 120 Ma and ending at 23 Ma (so that the oldest simulated “family” is always 120 Ma old and all families are older than Neogene). Analysis of these 50 x 198 clades found that only three datasets (6%) supported a pre-Cretaceous origin with P > 0.95 and even in these instances the probability of a Cretaceous origin is ≥0.04 (Fig. 1b; Extended Data Table 3). This is twenty times higher than the probability we estimated from the angiosperm fossil record, and three orders of magnitude higher than the probability estimated by Budd and colleagues in their simulation. These results show that even if angiosperms evolved under an improbable constant birth-death process starting in the early Cretaceous, our model would likely (P = 0.94) infer their origination time correctly.

Budd and colleagues suggest that the biasing impact of a small number of ancient angiosperm families with poor fossil records could be overcome by analysing the data at more inclusive taxonomic ranks. Orders are represented by more fossil samples than families and they show that BBB analyses of these data yield younger and more precise clade age estimates. However, precision does not equal accuracy and ordinal ages still favour a Jurassic origin of crown-angiosperms (with probability P = 0.59 and 95% CI of the root age 195–124 Ma; Extended Data Table 4). Their approach obscures the obvious signal in the raw fossil data-that lineages exhibit different sampling rates. To estimate among-lineage sampling rates it would be more appropriate to analyse the data at more exclusive taxonomic ranks (genus, species). Our family-rank analyses provide the most taxonomically inclusive level provided by the available fossil record, allowing for among-lineage variance in fossilization rates and avoiding lumping the data to produce specific results.

Budd and colleagues conjecture a pre-Cretaceous origin of crown-angiosperms, but not “*to the level implied by Silvestro et al*.”, reflecting a misreading of our results (^1^ Fig. 4), the uncertainties of which extend from early Cretaceous to late Permian for 1.5% of families. These uncertainties represent agnosticism as to the true time of divergence of these families. As such, our results are entirely compatible with those derived heuristically by Budd and colleagues^2^, and remain essentially unchanged even after excluding the early records of Papaveraceae and Triuridaceae, which they claim are problematic (Fig. 1c, d; Extended Data Table 5).

Budd and colleagues question the utility of the BBB model on the basis that its results are consistent, yet not identical, with a traditional Confidence Interval method^7^ applied to empirical data. Obtaining consistent results from different methods indicates that the signal in the data is strong. It is not a way to benchmark the performance of a model^8^.

In conclusion, Budd and colleagues’ concerns with the BBB model and its application to the angiosperm fossil record are unsound. The fossil record cannot be taken at face value since it always underestimates the true age of clades^9^. It cannot be interpreted heuristically based on generalized expectations of the diversification dynamics of stem-and crown-groups^10^, not least since these expectations are demonstrably violated by the known fossil record^11^. Interpretation of the fossil record requires modelling and the BBB model^5^ is a powerful and effective method for interpreting this important archive of evolutionary history.

## Authors’ contribution

DS performed the analyses. DS wrote the manuscript with contributions from CDB, PCJD, AA, WD, QZ, and YX.

---

**Extended Data Table 1**. Results from the analysis of 200 simulated clades with sampling probability set to 0. These datasets therefore only consist of the present diversity (column: N. extant sp.), without any fossil record. The table reports the true and estimated ages with 95% CI.

**Extended Data Table 2**. Results from the analysis of 198 simulated clades with origination time set to 120 Ma and default settings as implemented in the rootBBB program (github.com/dsilvestro/rootBBB). The table reports the true and estimated ages with 95% CI, number of extant species and number of simulated fossil occurrences.

**Extended Data Table 3**. Results from the analysis of 50 datasets each including 198 simulated clades. The ages of the simulated clades were drawn from a birth-death process with speciation rate set to 0.1 and no extinction (since all families in our empirical data are extant) with the age of the first clade set to 120 Ma and a minimum age boundary at 23 Ma. The table reports simulated and estimated ages for each clade in each of the 50 datasets along with an estimated probability that the crown age of the dataset is pre-Cretaceous (Prob root > 145).

**Extended Data Table 4**. Results from the re-analysis of angiosperm fossil record at the order level under the BBB model with time-increasing preservation rates. The table reports the number of modern species, number of fossil occurrences as well as the estimated age, initial sampling rate (*q*), slope of sampling increase through time (*a*) and log-variance of the Brownian bridge (σ^2^) and the respective 95% credible intervals.

**Extended Data Table 5**. Results from the re-analysis of angiosperm fossil record at the family level after removing the records considered as problematic by Budd et al.^2^ (resulting in the removal of the Triuridaceae family and of one occurrence within the Papaveraceae family) under the BBB model with time-increasing preservation rates. The table reports the number of modern species, number of fossil occurrences as well as the estimated age, initial sampling rate (*q*), slope of sampling increase through time (*a*) and log-variance of the Brownian bridge (σ^2^) and the respective 95% credible intervals.

## References

1 Silvestro, D. et al. Fossil data support a pre-Cretaceous origin of flowering plants. Nature Ecology & Evolution, doi:10.1038/s41559-020-01387-8 (2021).

2 Budd, G. E., Mann, R. P., Doyle, J. A., Coiro, M. & Hilton, J. Fossil data do not support a long pre-Cretaceous history of flowering plants. bioRxiv, doi: 10.1101/2021.02.16.431478 (2021).

3 Drummond, C. S., Eastwood, R. J., Miotto, S. T. & Hughes, C. E. Multiple continental radiations and correlates of diversification in Lupinus (Leguminosae): testing for key innovation with incomplete taxon sampling. Syst Biol 61, 443–460, doi:10.1093/sysbio/syr126 (2012).

4 Zanne, A. E. et al. Three keys to the radiation of angiosperms into freezing environments. Nature 506, 89–92, doi:10.1038/nature12872 (2014).

5 Silvestro, D., Cascales-Minana, B., Bacon, C. D. & Antonelli, A. Revisiting the origin and diversification of vascular plants through a comprehensive Bayesian analysis of the fossil record. The New phytologist 207, 425–436, doi:10.1111/nph.13247 (2015).

6 Magallón, S. & Castillo, A. Angiosperm diversification through time. Am. J. Bot. 96, 349–365, doi:10.3732/ajb.0800060 (2009).

7 Marshall, C. R. Confidence-intervals on stratigraphic ranges. Paleobiology 16, 1–10 (1990).

8 Sim, S. E., Easterbrook, S. & Holt, R. C. in 25th International Conference on Software Engineering (ICSE’03). (IEEE).

9 Donoghue, P. C. & Yang, Z. The evolution of methods for establishing evolutionary timescales. Philos Trans R Soc Lond B Biol Sci 371, 20160020, doi:10.1098/rstb.2016.0020 (2016).

10 Budd, G. E. & Mann, R. P. The dynamics of stem and crown groups. Science Advances 6, 1–10 (2020).

11 Jablonski, D. Lessons from the past: evolutionary impacts of mass extinctions. Proc Natl Acad Sci U S A 98, 5393–5398, doi:10.1073/pnas.101092598 (2001).

